# Serotonin stimulates proliferation of ionocytes via the 5-HT2A receptor in zebrafish larvae

**DOI:** 10.1101/2025.10.24.684416

**Authors:** Willamina M. MacDonald, Michael G. Jonz

**Author notes:** Author for correspondence: Michael G. Jonz, Ph.D, Department of Biology, University of Ottawa, 30 Marie Curie Pvt, Ottawa, ON K1N 6N5, Canada.

## Abstract

Osmoregulation is an essential process in all living organisms. For aquatic organisms, such as freshwater fishes whose natural environment is hyperosmotic, specialized cells, called ionocytes, are present in the skin during developmental stages and contribute to the maintenance of osmotic homeostasis. Such cells are known to proliferate in response to osmotic stress, but the molecular mechanism by which that process is regulated remains poorly characterized. In this study, using immunohistochemistry and confocal microscopy, we demonstrate that cutaneous ionocytes in developing zebrafish (*Danio rerio*) express serotonin 2A (5-HT2A) receptors by co-labeling with other known ionocyte markers, such as the Na^+^/K^+^-ATPase, Concanavalin A and Mitotracker. Furthermore, by quantifying ionocyte number through early stages of development, we implicate 5-HT2A receptors in initiating ionocyte proliferation. Exposure of zebrafish embryos and larvae to acidic pH, or exogenous 5-HT, increased the number of cutaneous ionocytes. The effects of both stimuli were abolished in the presence of the 5-HT2A receptor-specific antagonist, ketanserin. Moreover, activation of 5-HT2A receptors led to increased detection of ionocytes with phosphorylated extracellular signal-related kinase (ERK), a key regulator of cell division and differentiation linked with 5-HT2A. We used tetrabenazine, an inhibitor of vesicular monoamine transporter 2 (vmat2) and 5-HT storage, to deplete potential sources of 5-HT. Tetrabenazine treatment in fish exposed to acidic pH reduced ionocyte proliferation, implicating an endogenous source of 5-HT in regulation of ionocyte populations.These results demonstrate the importance of a pathway initiated by 5-HT2A activation that regulates ionocyte proliferation in developing zebrafish exposed to osmotic stress.

**SUMMARY STATEMENT:** Serotonin stimulates 5-HT2A receptors on ionocytes leading to proliferation upon acclimation to acidic environments in zebrafish larvae.

## INTRODUCTION

All living organisms must be equipped with mechanisms to maintain osmotic homeostasis. In the case of freshwater fishes, whose natural environment is hyperosmotic, osmoregulation includes excretion of dilute urine following reabsorption of ions in the kidney, and active absorption of ions through specialized cells called ionocytes (Evans et al., 2005). Ionocytes can be classified based on the ion transporters they express, many of which are analogous to transporters in renal tubular cells, the principal site of osmotic regulation in terrestrial vertebrates (Hwang and Chou, 2013). In zebrafish (*Danio rerio*), ionocytes are present in the gills of adults and the skin of developing fish. They can be divided into five populations: cells that are H^+^-ATPase-rich (HR) or Na^+^/K^+^-ATPase-rich (NaR), those that express the Na^+^/Cl^-^ cotransporter (NCC) or solute carrier 26 (SLC26), and K^+^-secreting (KS) cells (Guh et al., 2015). Ionocytes in the skin of developing zebrafish are especially convenient to study as they are directly exposed to the environment and are easily observed *in situ* with whole-mount preparations (Hwang and Chou, 2013).

In response to osmotic stress, ionocytes increase in number and in their expression of ion transporters. Cutaneous ionocytes of zebrafish larvae have previously been shown to proliferate with exposure to acidic (Horng et al., 2009; Shir-Mohammadi and Perry, 2020) and low Na^+^ environments (Shir-Mohammadi and Perry, 2020). It has also been demonstrated that individual HR ionocytes increase their H^+^-secreting function in response to acidic environments (Horng et al., 2009). The molecular mechanisms underlying these phenomena have yet to be fully characterized. Thus far, hormones such as prolactin (Breves et al., 2014) and isotocin (Chou et al., 2011) have been proposed as major regulators of ionocyte number. Furthermore, cortisol has been demonstrated to play a role in proliferation of gill ionocytes and an increase in Na^+^/K^+^-ATPase activity, at least partially by direct stimulation (Evans et al., 2005). With respect to the increase in function of single HR cells, adrenergic systems have been suggested to play a role. It has been demonstrated that HR cells express 𝛽-adrenergic receptors, and that Na^+^ uptake in these cells is altered by acute treatment with adrenergic agonists and antagonists (Kumai et al., 2012). These findings clearly demonstrate that regulation of ionocyte number and function depends on multiple factors that play a role in stress responses, raising the question of whether additional hormones or neurotransmitters that mediate stress responses in fishes are involved.

One such neurotransmitter is serotonin (5-hydroxytryptamine, 5-HT), which has been shown to become elevated in the brain of zebrafish exposed to hypoxic or osmotic stressors (Sreelekshmi et al., 2022). Increased 5-HT levels have also been observed in the brain (Boeck et al., 1996) and in the gills (Mustafayev and Mekhtiev, 2008) of other species of fish exposed to high salinity. An increase in 5-HT levels in the gill is especially interesting because ionocytes are located mainly in the gills of adult fishes, and are in close proximity to neuroepithelial cells (NECs) that retain and release 5-HT (Jonz and Nurse, 2006; Reed et al., 2025). Whether an increase in 5-HT levels occurs in the skin of zebrafish larvae exposed to osmotic stressors has not been demonstrated, though cutaneous, 5-HT-containing NECs are also present (Coccimiglio and Jonz, 2012) and are adjacent to cutaneous ionocytes (Dean et al., 2017). Given that acute treatment of zebrafish larvae with 5-HT does not lead to changes in Na^+^ uptake (Kumai et al., 2012), it is unlikely that 5-HT affects the expression or activity of ion transporters at the level of single ionocytes. However, it has yet to be investigated whether 5-HT might lead to proliferation of ionocytes.

5-HT has previously been demonstrated to be involved in regulation of cell number with effects on cell proliferation, maturation and apoptosis in various cell types and organisms including cells of the lung, kidney, and nervous system (Azmitia, 2001). Interestingly, it has been demonstrated in rat that 5-HT induces proliferation of renal mesangial cells (Takuwa et al., 1989) mediated by 5-HT2A receptors through pathway(s) involving extracellular signal-regulated kinases (ERKs) (Göőz et al., 2006; Greene et al., 2000; Grewal et al., 1999).

The present study employed immunohistochemistry and confocal microscopy to reveal that 5-HT2A receptors are found in cutaneous ionocytes in zebrafish larvae. We hypothesized that 5-HT2A receptors facilitate regulation of ionocyte number in response to osmotic stress. To test this hypothesis, we first aimed to demonstrate that 5-HT2A-immunolabeling co-localized with known markers of multiple ionocyte subtypes. We then aimed to demonstrate a role for 5-HT, via the 5-HT2A receptor, in regulating ionocyte number. To do so, we quantified the number of ionocytes under different conditions, including acid acclimation and treatment with 5-HT2A-specific agonists and antagonists. Furthermore, we demonstrate that the source of 5-HT is a cell type that utilizes vesicular monoamine transporter 2 (vmat2), potentially NECs. Finally, using an antibody raised against phosphorylated ERK, we provide evidence that the pathway by which the 5-HT2A receptor leads to proliferation of ionocytes involves ERK and that phosphorylated ERK is present only in 5-HT2A-positive ionocytes that have been stimulated by 5-HT.

The results highlight a role for 5-HT, via 5-HT2A receptors, in initiating proliferation of ionocytes in response to low pH environments. This novel role for 5-HT in zebrafish larvae contributes to our evolving understanding of the mechanisms by which zebrafish respond to osmotic stressors at early developmental stages. These findings may have implications in understanding how fish respond to osmotic challenges in the environment and in aquaculture, as well as potential applications for zebrafish as a model organism to enhance our understanding of analogous cell types and pathways in mammals.

## MATERIALS AND METHODS

### Animal care

Zebrafish were maintained at the Laboratory for the Physiology and Genetics of Aquatic Organisms, University of Ottawa, at 28°C on a 14:10-h light:dark cycle (Westerfield, 2007). Water entering the facility from the city of Ottawa was dechloraminated and aerated. System water pH was buffered using Proline Sodium Bicarbonate (NaHCO_3_) following manufacturer’s directions (cat. no. PC12, Pentair Aquatic Eco-Systems, Inc., Apopka, FL, USA). Wild-type male and female zebrafish were used for all experiments. Embryos were bred from 12-month adults using standard breeding techniques (Westerfield, 2007) and transferred to 150-mm Petri dishes containing buffered system water and 3.0×10^-5^ % methylene blue (unless otherwise stated) at 0-2 hours post-fertilization (hpf). Dishes were kept in an incubator at 28.5°C and media was replaced daily. When the desired age (2-7 days postfertilization, dpf) was reached, larvae were euthanized in 1 mg/mL MS-222 buffered with 0.5 mg/mL NaHCO_3_ at 4°C. All animal use procedures were carried out according to institutional guidelines and protocol BL-3666 in accordance with the Canadian Council on Animal Care.

### Immunohistochemistry

Euthanized zebrafish larvae were processed for whole-mount immunohistochemistry according to previously established protocols (Coccimiglio and Jonz, 2012; Jonz and Nurse, 2005). In brief, samples were processed with: (1) fixation, (2) permeabilization, (3) incubation in primary antibodies, and (4) incubation in secondary antibodies. Between each step, the larvae were rinsed 3 times, for 3 min in phosphate buffered solution (PBS) containing (mM): NaCl 137, Na_2_HPO_4_ 15.2, KCl 2.7, and KH_2_PO_4_ 1.5 at pH 7.8. Fixation was done in 4% paraformaldehyde (PFA) in PBS at 4°C overnight. Permeabilization was then accomplished by incubating the samples in 2% Triton X-100 (TX-100) in PBS at 4°C for 48 h. Primary and secondary antibodies were prepared by dilution in PBS to 1:100. Fish were incubated in primary antibodies at 4°C for 24 h. Samples were then incubated in secondary antibodies for 1 h at room temperature in a dark chamber. In double-labeling experiments, antibodies were used in combination. Finally, samples were mounted on glass slides in ProLong Diamond Antifade Mountant (cat. no. P36970; Thermo Fisher Scientific, Waltham, MA, USA).

To demonstrate that 5-HT2A receptors were present in ionocytes, a 5-HT2A receptor antibody was used. The polyclonal 5-HT2A antibody (RRID_AB2807092, cat. no. PA5-95288, lot no. 79627209 and no. ZG4406664A, Thermo Fisher Scientific) was raised in rabbit against a synthetic peptide corresponding to the C-terminus of human 5-HT2A receptor, specifically amino acids 400 to 431 (manufacturer specifications). This corresponds to residues 265-296 of the zebrafish 5-HT2A receptor (Fig. S1). To recognize the 5-HT2A antibody, two different secondary antibodies were used depending on the experiment. In figures where 5-HT2A labeling is magenta, a polyclonal anti-rabbit Alexa 594-conjugated antibody (cat. no. A11012, lot no. 3112802 and 2616076, Thermo Fisher Scientific) raised in goat was used. In figures where 5-HT2A labeling is green, a polyclonal anti-rabbit FITC-conjugated antibody (cat. no. 111-095-003, lot no. 104432, Jackson ImmunoResearch) raised in goat was used. Since the 5-HT2A antibody has not previously been used to label ionocytes in zebrafish larvae, we performed a negative control. To do so, we synthesized a custom peptide corresponding to the antigen sequence: KENKKPLQLILVNTIPALAYKSSQLQMGQKKN (lot no. ABc10124, ABclonal Science, Woburn, MA, USA). Primary antibodies were pre-incubated with the peptide at a concentration 5 times greater than that of the antibody at room temperature for 1 h before adding tissue. No immunolabeling was seen in pre-adsorbed samples, demonstrating that the antibody was not binding via off-target effects.

To provide evidence that the observed 5-HT2A-positive cells were ionocytes, ionocytes were co-labeled with well-established markers previously used to identify cutaneous ionocytes in zebrafish larvae. Such markers include an antibody raised against the 𝛼5 subunit of Na^+^/K^+^-ATPase, a known marker of NaR ionocytes (Hwang and Chou, 2013; Jonz and Nurse, 2006), Concanavalin A (ConA), a known marker of HR ionocytes (Esaki et al., 2007; Horng et al., 2009; Kumai et al., 2012; Kwong and Perry, 2016), and Mitotracker, a known marker of mitochondrion-rich cells (MRCs) including many ionocyte subtypes, namely NaR, HR, and NCC ionocytes (Esaki et al., 2007; Kumai et al., 2012; Kwong and Perry, 2016). The monoclonal α5 primary antibody (RRID: AB_2166869, Developmental Studies Hybridoma Bank, University of Iowa, USA) was raised in mouse against Na^+^/K^+^-ATPase from chicken kidney and recognizes a cytosolic epitope on the α subunit of Na^+^/K^+^-ATPase (manufacturer specifications). It was recognized by a polyclonal Alexa 488-conjugated anti-mouse secondary antibody (cat. no. A11029, lot no. 57465A, Thermo Fisher Scientific) raised in goat. ConA and Mitotracker are both conjugated to fluorophores. For details, see the *Chemical exposures* section.

To demonstrate activation of a pathway involving ERK, an antibody raised against phosphorylated ERK (p-ERK) was employed. Monoclonal anti-p-ERK (RRID: AB_2572926, cat. no. 14-9109-82, clone no. MILAN8R, lot no. 2643096, Invitrogen, Burlington, ON, Canada) was raised in mouse. Target specificity was previously validated in zebrafish larvae, where the p-ERK antibody was used as a biomarker for neuronal activation (Spadacini, 2024). In the present study, p-ERK labelling in the absence of other antibodies was done to test for cross-reactivity. Anti-pERK demonstrated the same labelling patterns with and without co-labelling with 5-HT2A. Anti-p-ERK was recognized with a polyclonal Alexa 594-conjugated anti-mouse secondary antibody (cat. no. A11005, lot no. 2043369, Invitrogen) raised in goat.

### Acid acclimation

Developing zebrafish were exposed to low pH to induce ionocyte proliferation. For these experiments, embryos were transferred to acidic medium immediately after collection. Acidic medium was composed of system water supplemented with 300 𝜇M 2-(N-morpholino) ethanesulfonic acid hydrate (MES) (cat. no. M8250; Sigma-Aldrich, Oakville, ON, Canada), adjusted to pH 4 with HCl, as done by Horng et al. (2009). Zebrafish were exposed immediately following embryo collection (0-2 hpf) for 2 to 7 days and kept in a 28.5°C incubator. Dishes contained no more than 60 embryos or larvae. Controls were treated in the same way but maintained in buffered system water.

Initial acid acclimation was done at developmental stages from 2 to 7 dpf to indicate which length of acid exposure and developmental stage would allow observation of robust changes in ionocyte number. Such experiments (Fig. 5) were done as one set of exposures with 5 fish sampled at each developmental stage and processed for immunohistochemistry. Following this initial experiment, all larvae were exposed to stimuli for 6 days (from 0 to 6 dpf).

### Chemical exposures

For experiments using Mitotracker to label ionocytes, Mitotracker Red CMXRos (cat. no. M7512; Invitrogen) was first dissolved in dimethyl sulfoxide (DMSO) (cat. no. D8418; Sigma) to a concentration of 1 mM according to the manufacturer’s directions. The stock solution was then diluted in extracellular solution (ECS) containing (mM): 120 NaCl, 5 KCl, 2.5 CaCl_2_, 2 MgCl_2_, 10 HEPES, 10 glucose at pH 7.8 to a final concentration of 300 nM Mitotracker. Mitotracker exposures were performed in ECS as opposed to embryo medium to avoid potential adverse effects of methylene blue on mitochondria or uptake of Mitotracker dye. Methylene blue has been shown to interact with mitochondria by multiple mechanisms (Klosowski et al., 2020). For experiments in which HR ionocytes were labeled with ConA, Concanavalin A-Alexa Fluor 488 conjugate (cat. no. C11252; Invitrogen) stock solutions were prepared to a final concentration of 5 mg/mL in 0.1 M NaCO_3_ (pH 8.3) according to the manufacturer’s instructions. The ConA stock was then diluted in ECS to a final concentration of 50 𝜇g/mL. Fish treated with Mitotracker or ConA were incubated in the desired dye for 30 min at 28.5°C before euthanasia and immunohistochemistry (described above).

To implicate the serotonergic system in ionocyte proliferation, zebrafish were exposed to 5-HT and the number of ionocytes on the trunk was quantified. To demonstrate that the effect of 5-HT was mediated by the 5-HT2A receptor, a 5-HT2A receptor-specific antagonist, ketanserin, was used. Finally, to begin to identify the source of 5-HT, a vesicular monoamine transporter 2 (vmat2) inhibitor, tetrabenazine, was used. Tetrabenazine was used because NECs (a known source of 5-HT in the skin) use vmat2 to load 5-HT into vesicles preceding release upon stimulation. For experiments in which zebrafish larvae were treated with 5-HT (cat. no.B21263.03; Thermofisher), ketanserin (cat. no. S006; Sigma), or tetrabenazine (cat. no. T2952; Sigma), the appropriate medium (control, or pH 4) was supplemented with 100 𝜇M of the drug of interest. The embryos were transferred to the appropriate medium immediately after collection (0-2 hpf) and exposed for 6 days (until 6 dpf). Solutions were prepared fresh daily when medium was changed. For solubility, ketanserin and tetrabenazine were first prepared as 100 mM stock solutions in DMSO. The final concentration of DMSO in the medium was 0.1%, and controls were performed with 0.1% DMSO, where appropriate. For fish treated with both ketanserin and 5-HT, fish were pre-incubated with ketanserin for 1 h before the addition of 5-HT daily. There was no change in pH with the addition of 5-HT, ketanserin, or tetrabenazine.

### Imaging and analysis

Images were taken with an upright Olympus FV1000 BX61 LSM confocal microscope with a H101A ProScan platform equipped with continuous wave laser lines at 405, 488 and 559 nm. Images were viewed and captured with Fluoview software (Olypmus, Richmond Hill, ON, Canada). Images were taken in optical sections and rendered as stacks for rotation. Image processing was done in FIJI (Schindelin et al., 2012). Signals from red (Alexa 594, or Mitotracker) fluorophores were converted to magenta.

For experiments in which cell number was quantified, images were taken at 40× magnification. Cells were counted manually using cell counter in FIJI in three groups: cells labeled with Mitotracker only (magenta), cells labeled with 5-HT2A only (green), and co-labeled cells (white) for the quantification of ionocyte number. For quantification of p-ERK labeling, cells were counted in two groups: cells labeled with 5-HT2A only, and cells labeled with both 5-HT2A and p-ERK. Only cells on the trunk were counted. The fin fold contained very few ionocytes and was excluded from analysis. A depiction of the boundary set for quantification of cell number on the trunk is shown with a dotted line in Figure 5A5 and A6. Only ionocytes were counted, and were identified based on location and morphology. The only other 5-HT2A-labeled cells were neuromast cells (also indicated in Figure 5A5 and A6), which were easily identified and excluded from further analysis.

Statistical analysis was performed with Prism v.9.5.1 (GraphPad Software Inc., San Diego, CA, USA). For comparisons of cell number over the course of development and for the number of p-ERK positive cells, a Multiple Mann-Whitney test with two-stage linear set-up procedure of Benjamin, Krieger and Yekutieli was used. For comparisons of cell number at 6 dpf under different conditions a Kruskal-Wallis test with Dunn’s multiple comparison test was used. For comparison of the percentage of p-ERK-positive cells, a Mann-Whitney test was used. All data is presented as means ± standard deviation (s.d.). Sample size (N) refers to the number of fish and p < 0.05 for all analyses.

## RESULTS

### 5-HT2A immunohistochemical labeling co-localized with known markers of ionocytes

Preliminary observation of 5-HT2A labeling in the skin of zebrafish larvae demonstrated ionocyte-like labeling patterns. At 3 dpf, the observed 5-HT2A-positive cells were distributed on the yolk sac, along the trunk, and began to appear on the head (Fig. 1). Therefore, we sought to co-localize the 5-HT2A antibody with known markers of ionocytes. The 5-HT2A antibody localized to the same cells as 𝛼5, ConA and Mitotracker. 5-HT2A and 𝛼5 labeling was generally confined to the plasma membrane; whereas Mitotracker was found in the cytoplasm, and ConA was localized to apical pits of HR cells, as previously described (Horng et al., 2009). All observed cells labeled with 𝛼5 (Fig. 2) or ConA (Fig. 3) were also labeled with 5-HT2A (N=25 and N=15, respectively). Similarly, 97% (N=160) of cells labeled with Mitotracker also contained 5-HT2A (Fig. 4). Mitotracker, the most broad-spectrum marker of ionocytes employed, labeled the majority, approximately 76% (N=160), of the observed 5-HT2A-positive cells. The remaining 24% of 5-HT2A-positive cells not labeled with Mitotracker were likely ionocytes, based on their morphology (Fig. 4), distribution, and increase in number following acid exposure (Fig. 5).

**Figure 1.**
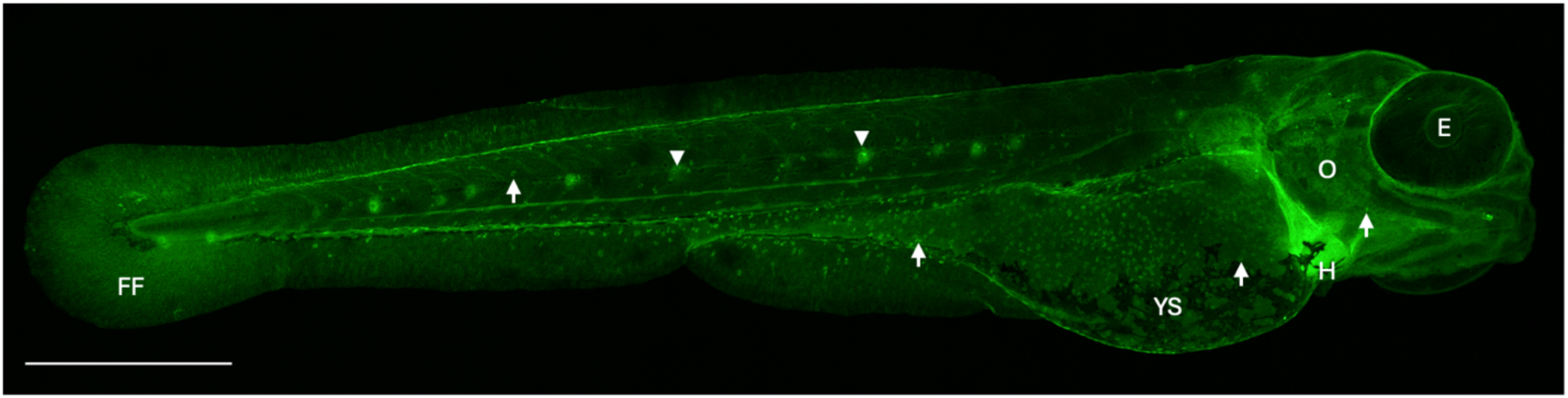
Overview of 5-HT2A labeling in zebrafish larvae. Representative confocal image of anti-5-HT2A labeling (green) in a zebrafish larva at 3 days postfertilization (dpf). 5-HT2A immunohistochemical labeling consistently demonstrated this pattern in >180 fish between 2 and 7 dpf, alone and in combination with other markers. Images were taken at 10× magnification in three segments, which were aligned to create a montage of the entire fish. Scale bar is 500 μm. Arrows point to examples of cells characterized as ionocytes, and arrowheads to neuromast cells. YS, yolk sac; H, heart; E, eye; O, operculum; FF, fin fold.

**Figure 2.**
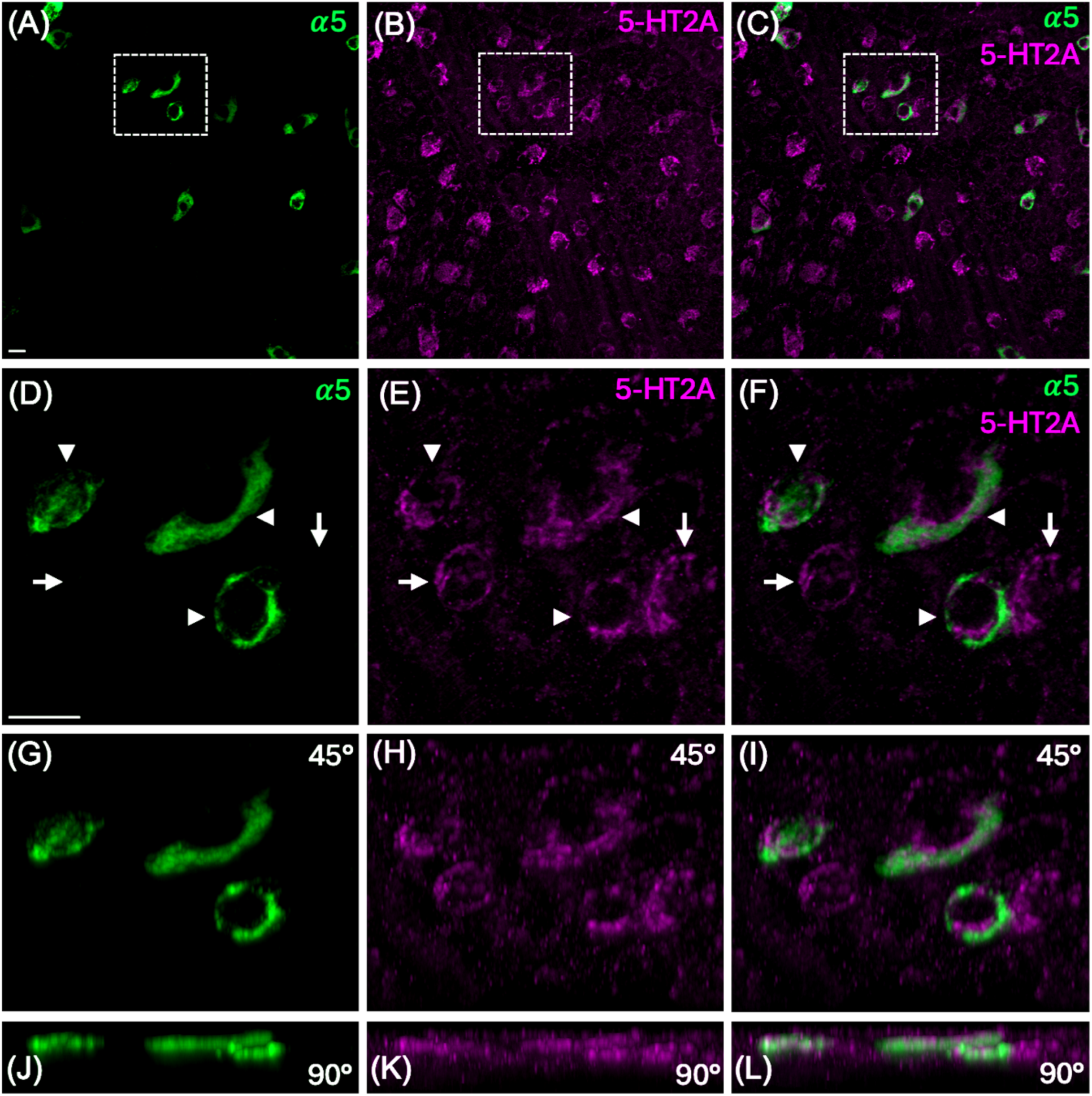
NaK (𝜶5)-positive ionocytes labeled positive for 5-HT2A. Representative confocal images of 𝛼5 (green) and 5-HT2A (magenta) labeling in the yolk sac of zebrafish at 3 days postfertilization (dpf). 5-HT2A and 𝛼5 antibodies consistently demonstrated this pattern in 25 zebrafish at 3 dpf across four rounds of immunohistochemistry. (A-C) Overview of 𝛼5 and 5-HT2A labeling. (D-L) Enlarged images of co-labeled cells (arrowheads) and cells positive for 5-HT2A only (arrows). (G-I, and J-L) Images from above rotated back 45° and 90°, demonstrating that 𝛼5 and 5-HT2A labeling were present in the same plane and were localized to the same cells. Scale bars are 10 μm. Scale bar in (A) applies to (A-C) and in (D) applies to (D-L). 𝛼5, Na^+^/K^+^-ATPase 𝛼 subunit antibody; 5-HT2A, serotonin 2A receptor.

**Figure 3.**
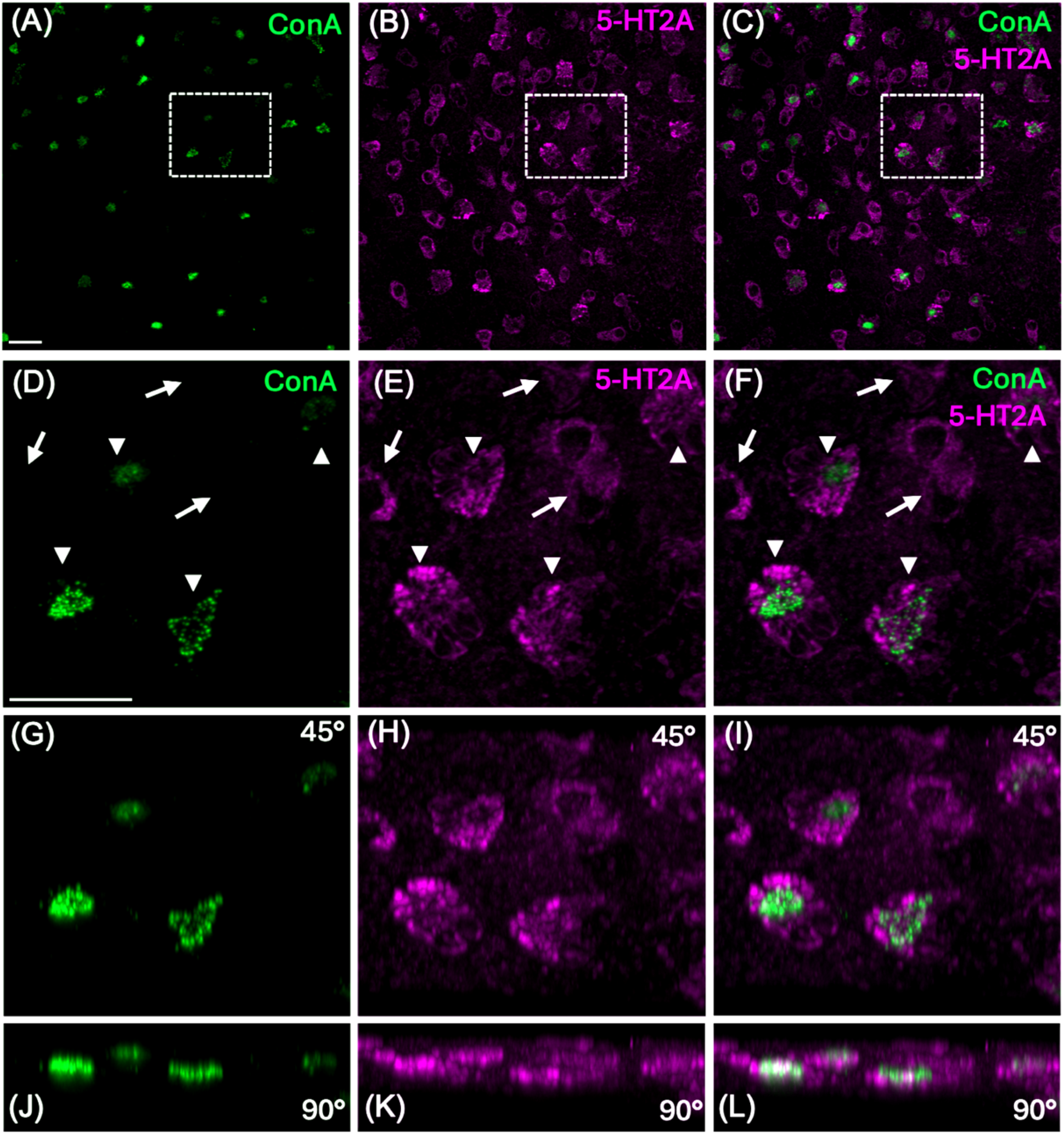
HR (ConA-positive) ionocytes were positive for 5-HT2A. Representative confocal images of ConA (green) and 5-HT2A (magenta) labeling on the yolk sac of zebrafish at 3 days postfertilization (dpf). The 5-HT2A antibody and ConA consistently demonstrated this pattern in 15 zebrafish at 3 dpf across two rounds of immunohistochemistry. (A-C) Overview of ConA and 5-HT2A labeling. (D-L) Enlarged image of co-labeled cells (arrowheads) and cells positive for 5-HT2A only (arrows). (G-I, and J-L) Images rotated back 45° and 90°, demonstrating ConA and 5-HT2A labeling were present in the same plane and were localized to the same cells. Scale bars are 20 μm. Scale bar in (A) applies to (A-C) and in (D) applies to (D-L). ConA, Concanavalin A; 5-HT2A, serotonin 2A receptor.

**Figure 4.**
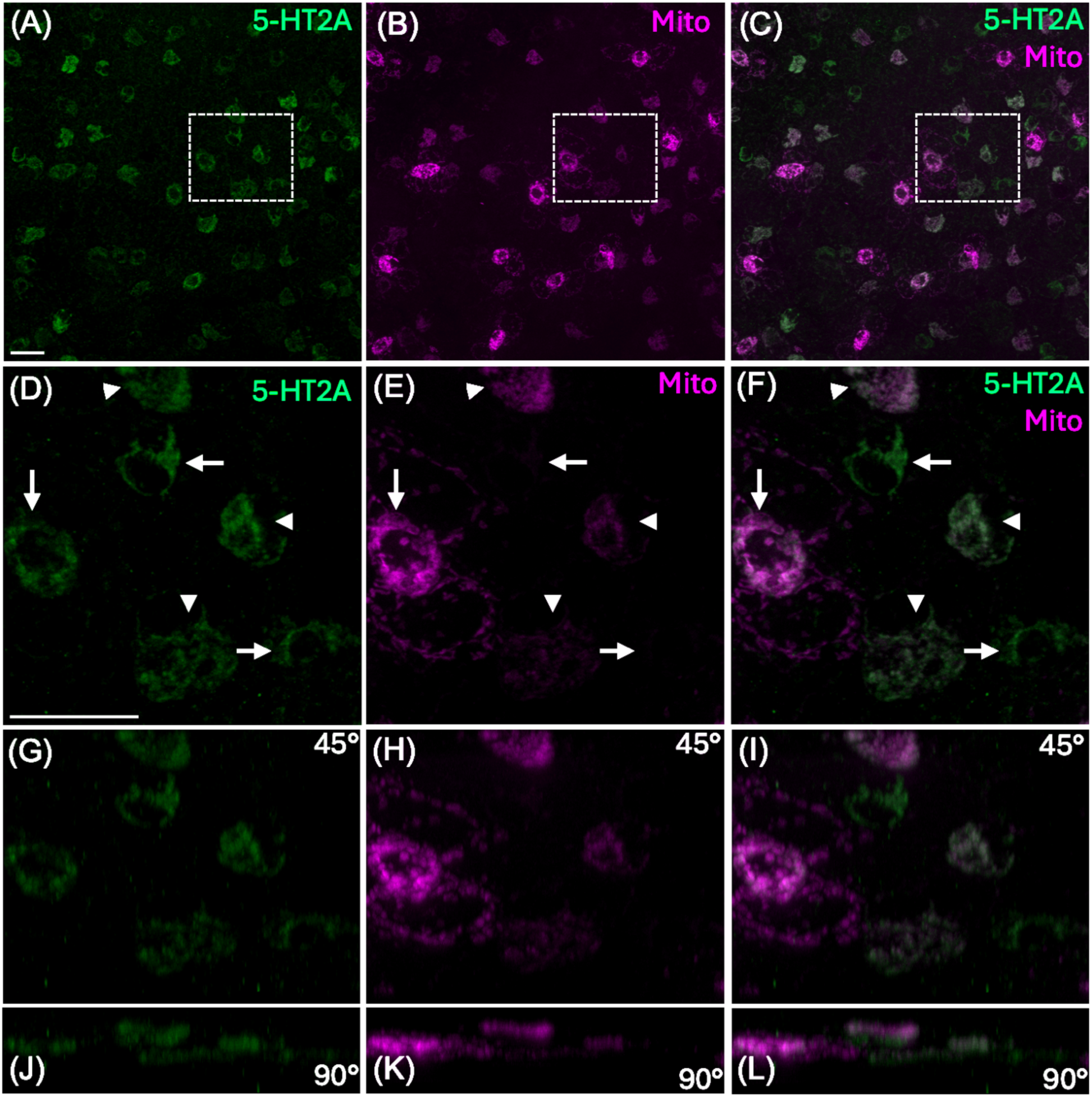
Mitochondrion-rich (Mitotracker positive) ionocytes were labeled by 5-HT2A antibodies. Confocal images of Mitotracker (magenta) and 5-HT2A (green) labeling on the yolk sac of zebrafish larvae at 3 days postfertilization (dpf). The 5-HT2A antibody and Mitotracker consistently demonstrated this pattern in 160 zebrafish between 2 and 7 dpf across multiple rounds of immunohistochemistry and different treatment conditions. (A-C) Overview of Mitotracker and 5-HT2A labeling. (D-L) Enlarged image of co-labeled cells (arrowheads) and cells positive for 5-HT2A only (arrows). (G-I, and J-L) Images rotated back 45° and 90°, demonstrating Mitotracker and 5-HT2A labeling in the same plane and localized to the same cells. Scale bars are 20 μm. Scale bar in (A) applies to (A-C) and in (D) applies to (D-L). 5-HT2A, serotonin 2A receptor; Mito, Mitotracker.

### 5-HT2A-positive ionocytes increased in number with exposure of zebrafish larvae to acidic medium

To elicit ionocyte proliferation, we treated zebrafish larvae with acidic medium (Fig. 5). A significant difference (p<0.05, N=5) in the total number of 5-HT2A-positive ionocytes during development was observed on the head (Fig. 5C) and trunk (Fig. 5E), but not on the yolk sac (Fig. 5D). This was the case for 5-HT2A-positive ionocytes co-labeled with Mitotracker (Fig. 5H,K) as well as ionocytes that were 5-HT2A-positive but Mitotracker negative (Fig. 5G,J) on the head and trunk. Cells that were positive for Mitotracker only were very few in number and did not demonstrate a significant change with acid exposure at any developmental stage (Fig. 5F,I). Given that they did not respond to treatment and did not contain 5-HT2A, these cells were not quantified in the remaining experiments. Compared to other regions, ionocytes on the trunk were best labeled and most evenly distributed, so only the trunk was used to quantify the change in ionocyte number in subsequent experiments. In addition, fish demonstrated a robust change in ionocyte number by 6 dpf, therefore this developmental stage was used in subsequent experiments.

**Figure 5.**
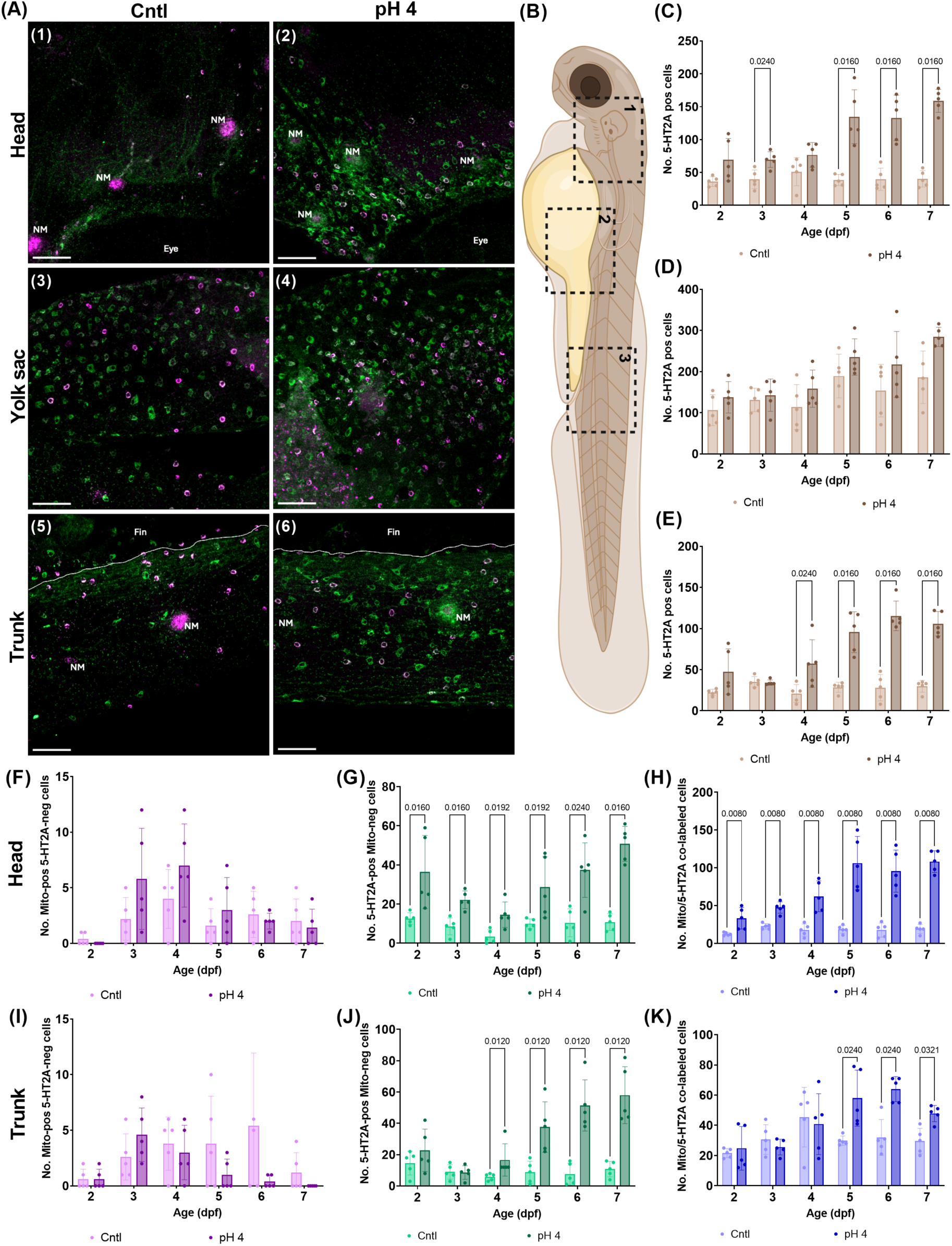
The number of 5-HT2A-positive cells on the head and trunk increased with acid exposure. (A) Representative confocal images of Mitotracker (magenta) and 5-HT2A (green) labeling on the head (1,2), yolk sac (3,4) and trunk (5,6) of zebrafish larvae at 7 days postfertilization (dpf) raised in control medium (1,3,5) or under pH 4 conditions (2,4,6). Fish were placed in the appropriate medium at approximately 1 h postfertilization, and 5 fish from each treatment group were collected and processed for immunohistochemistry daily from 2 to 7 dpf. Scale bars 50 𝜇m. (B) Area imaged for the head (1), yolk sac (2) and trunk (3). (C-E) The total number of ionocytes with acid treatment compared to the control on the head (C), yolk sac (D) and trunk (E). (F-H) Cells labeled with Mitotracker only (F), 5-HT2A only (G) and co-labeled with 5-HT2A and Mitotracker (H) on the head. (I-K) Cells labeled with Mitotracker only (I), 5-HT2A only (J) and co-labeled with 5-HT2A and Mitotracker (K) on the trunk. Data analyzed using a Multiple Mann-Whitney test (two-tailed) with two-stage linear set-up procedure of Benjamin, Krieger and Yekutieli. p values are shown on the graph for groups that are significantly different (p<0.05, N=5 for each group at a given age). Average values represented as mean±s.d. NM, neuromast; E, eye; FF, dorsolateral fin fold; 5-HT2A, serotonin 2A receptor; Cntl, control; pos, positive; neg, negative.

### The number of 5-HT2A-positive ionocytes increased with exposure to 5-HT, but not with the addition of ketanserin

Following the identification of 5-HT2A in proliferating ionocytes, we sought to investigate a role for the 5-HT2A receptor in increasing ionocyte number. Considering this hypothesis, we first exposed zebrafish larvae to the agonist, 5-HT, for 6 days (from 0-6 dpf). Following 5-HT exposure, the number of 5-HT2A-positive ionocytes was significantly increased (p<0.05, N=10), regardless of whether they were co-labeled with Mitotracker, as was the case for ionocyte number in pH 4-treated fish (Fig. 6B,C).

**Figure 6.**
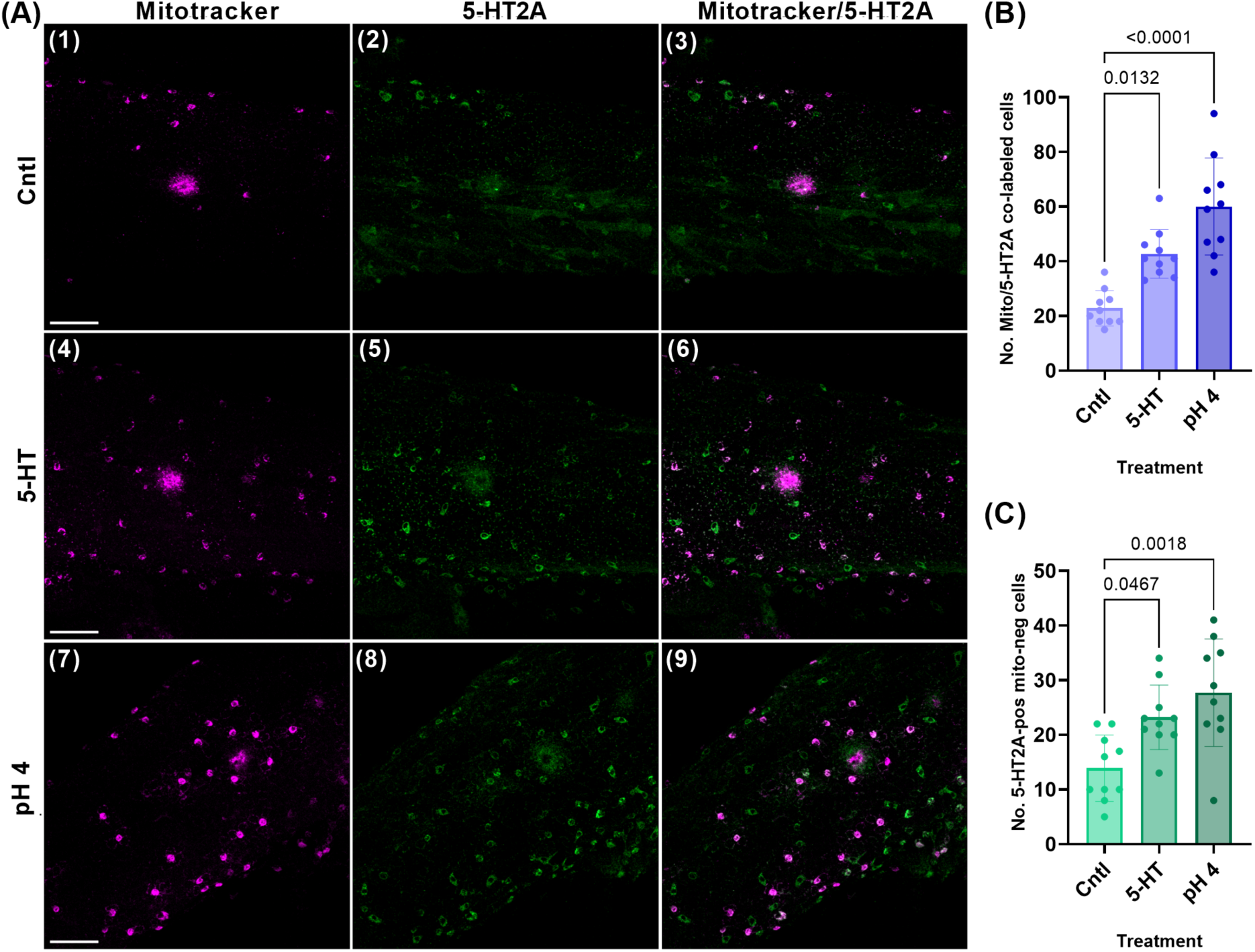
5-HT exposure resulted in an increased number of 5-HT2A-positive ionocytes. (A) Representative confocal images of Mitotracker (magenta) labeling (1,4,7), 5-HT2A (green) labeling (2,5,8), and Mitotracker/5-HT2A co-labeling (white) (3,6,9) on the trunk of zebrafish larvae at 6 days postfertilization (dpf) in control medium (1-3), medium supplemented with 100 𝜇M 5-HT (4-6), and at pH 4 (7-9). Fish were placed in the corresponding medium at approximately 1 h postfertilization and treated for 6 days (from 0 to 6 dpf). 10 fish from each of the three treatment groups were treated and processed for immunohistochemistry. Scale bars in (A) are 50 𝜇m and apply to all panels in that row. (B) The number of ionocytes co-labeled with Mitotracker and 5-HT2A. (C) The number of ionocytes labeled with 5-HT2A, but not with Mitotracker. Data analyzed using a Kruskal-Wallis test (two-tailed) with Dunn’s multiple comparison test. p values are shown on the graph for groups that are significantly different (p<0.05, N=10 for each group). Average values represented as mean±s.d. 5-HT, serotonin; 5-HT2A, serotonin 2A receptor; Cntl, control; Mito, Mitotracker; pos, positive; neg, negative.

In a subsequent round of experiments, we demonstrated that 5-HT was acting specifically through the 5-HT2A receptor to cause the increase in number of ionocytes. We pre-exposed zebrafish larvae to the specific 5-HT2A antagonist, ketanserin, before exposing to 5-HT. When fish were pre-exposed to ketanserin, the effect of 5-HT on ionocyte number was abolished, while 5-HT treatment in the absence of ketanserin continued to have a significant effect (p<0.05, N=10) (Fig. 7B,C). These experiments included solutions supplemented with 0.1% DMSO, since it was required to dissolve ketanserin. DMSO itself had no effect on ionocyte number (Fig. S2).

**Figure 7.**
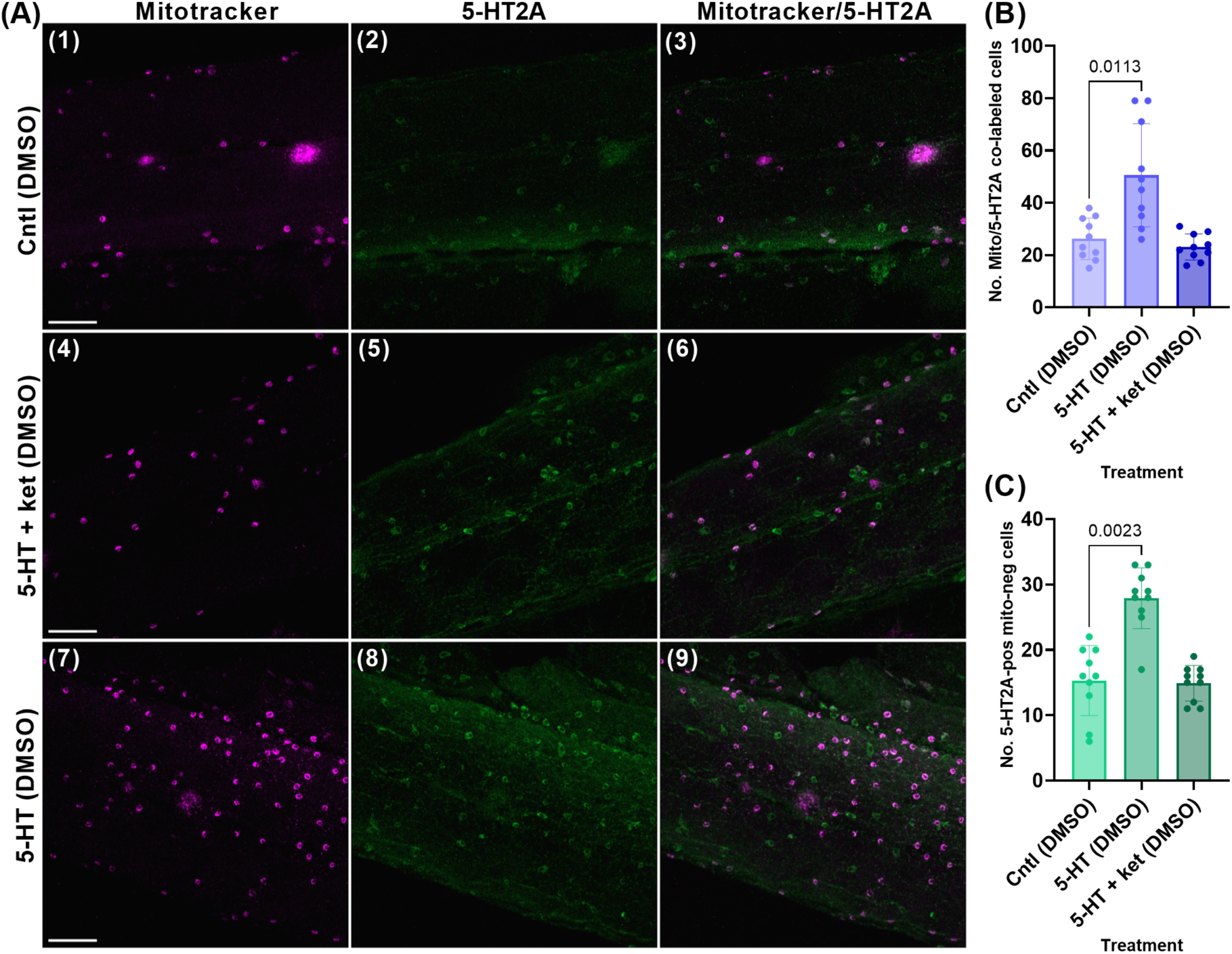
Pre-treatment with ketanserin inhibited the increase in ionocyte number elicited by 5-HT. (A) Representative confocal images of Mitotracker (magenta) labeling (1,4,7), 5-HT2A (green) labeling (2,5,8), and Mitotracker/5-HT2A co-labeling (white) (3,6,9) on the trunk of zebrafish larvae at 6 days postfertilization (dpf) under the following conditions: in control medium supplemented with 0.1% DMSO (1-3), in medium supplemented with 100 𝜇M 5-HT and 0.1% DMSO (4-6), and in medium with 100 𝜇M 5-HT following pre-treatment with 100 𝜇M ketanserin supplemented with 0.1% DMSO (7-9). Fish were placed in the corresponding medium at approximately 1 h postfertilization and treated for 6 days (from 0 to 6 dpf). 10 fish from each of the three treatment groups were treated and processed for immunohistochemistry. Scale bars in (A) are 50 𝜇m and apply to all panels in that row. (B) The number of ionocytes co-labeled with Mitotracker and 5-HT2A. (C) The number of ionocytes labeled with 5-HT2A, but not Mitotracker. Data analyzed using a Kruskal-Wallis test (two-tailed) with Dunn’s multiple comparison test. p values are shown on the graph for groups that are significantly different (p>0.05, N=10 for each group). Average values represented as mean±s.d. 5-HT, serotonin; 5-HT2A, serotonin 2A receptor; ket, ketanserin; DMSO, dimethyl sulfoxide; Cntl, control; Mito, Mitotracker; pos, positive; neg, negative.

### Treatment with 5-HT resulted in phosphorylation of ERK in ionocytes, which was reduced when fish were pre-treated with ketanserin

Given that ERK is commonly involved in signaling cascades initiated by the 5-HT2A receptor, we hypothesized that treatment with 5-HT would result in phosphorylation of ERK in 5-HT2A-positive ionocytes. We used an antibody raised against p-ERK to demonstrate activation of a pathway in ionocytes upon stimulation of 5-HT2A with 5-HT. Fish were processed for immunohistochemistry using anti-p-ERK and anti-5-HT2A. We quantified the number of p-ERK-positive and negative ionocytes in each fish and, subsequently, the percentage of 5-HT2A-positive ionocytes co-labeled with p-ERK (Fig. 8). In unstimulated controls, no clear p-ERK labeling was observed in any ionocytes. However, in 5-HT and ketanserin-treated samples, p-ERK-positive cells were observed (Fig. 8). In 5-HT-treated samples, the number of p-ERK-positive ionocytes was significantly greater (p<0.0001, N=10) than the number of p-ERK-negative ionocytes (Fig. 8B). In fish pre-treated with ketanserin, the number of p-ERK-positive ionocytes was significantly less (p<0.0001, N=10) than the number of p-ERK negative ionocytes (Fig. 8B). Furthermore, in 5-HT-treated samples, an average of about 74% of 5-HT2A-labeled ionocytes were p-ERK positive, which was significantly reduced (p<0.0001, N=10) to an average of about 23%, when fish were pretreated with ketanserin (Fig. 8C).

**Figure 8.**
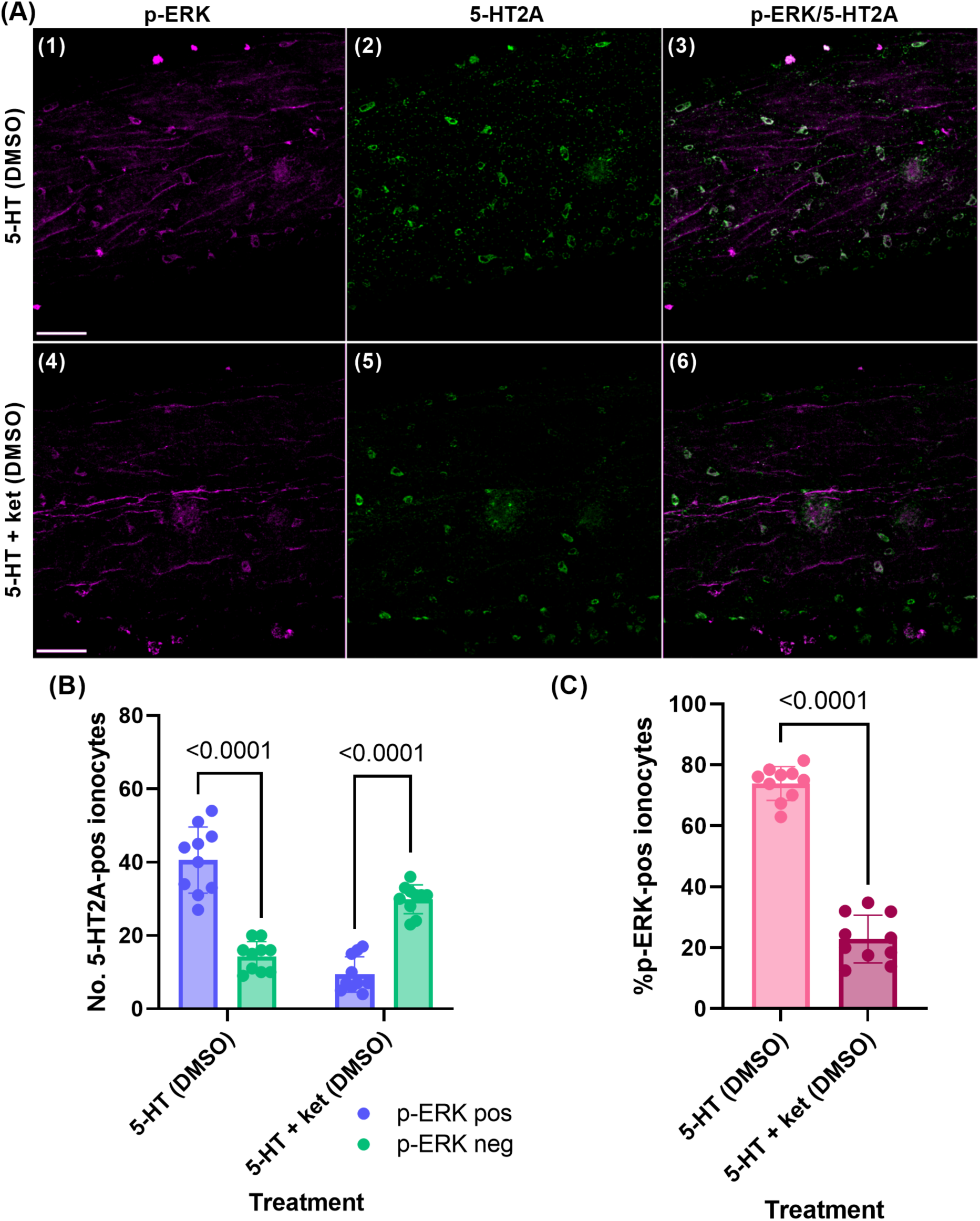
A higher percentage of ionocytes contained p-ERK when larvae were treated with 5-HT, than when pre-treated with ketanserin. (A) Representative confocal images of p-ERK (magenta) labeling (1,4), 5-HT2A (green) labeling (2,5), and p-ERK/5-HT2A co-labeling (white) (3,6) on the trunk of zebrafish larvae at 6 days postfertilization (dpf) under the following conditions: 100 𝜇M 5-HT supplemented with 0.1% DMSO (1-3); and 100 𝜇M 5-HT pre-treated with 100 𝜇M ketanserin supplemented with 0.1% DMSO (4-6). Fish were placed in the corresponding medium at approximately 1 h postfertilization and treated for 6 days (from 0 to 6 dpf). 10 fish from each of the treatment groups were treated and processed for immunohistochemistry. Scale bars in A1 and A4 are 50 𝜇m and apply to all panels in that row. (B) The number of p-ERK-positive ionocytes compared to the number of p-ERK-negative ionocytes in each treatment group. (C) The percentage of p-ERK-labeled 5-HT2A-positive ionocytes in each treatment group. Data in (B) were analyzed using a Multiple Mann-Whitney test (two-tailed) with two-stage linear set-up procedure of Benjamin, Krieger and Yekutieli. Data in (C) were analyzed using a Mann-Whitney test (two-tailed). p values are shown on the graph for groups that are significantly different (p<0.05, N=10 for each group). Average values represented as mean±s.d. 5-HT, serotonin; 5-HT2A, serotonin 2A receptor; p-ERK, phosphorylated extracellular signal-regulated kinase; ket, ketanserin; pos, positive; neg, negative.

### Ketanserin and tetrabenazine inhibited the observed increase in ionocyte number in acidic medium

Given that ketanserin inhibited the increase in ionocyte number observed in the presence of 5-HT, we next sought to investigate whether it could also inhibit the increase in ionocyte number with acclimation to an acidic environment (Fig. 9). Following exposure to acidic medium supplemented with ketanserin from 0-6 dpf, no significant increase (p>0.05, N=10) in ionocyte number was observed compared to the negative control (Fig. 9B,C).

**Figure 9.**
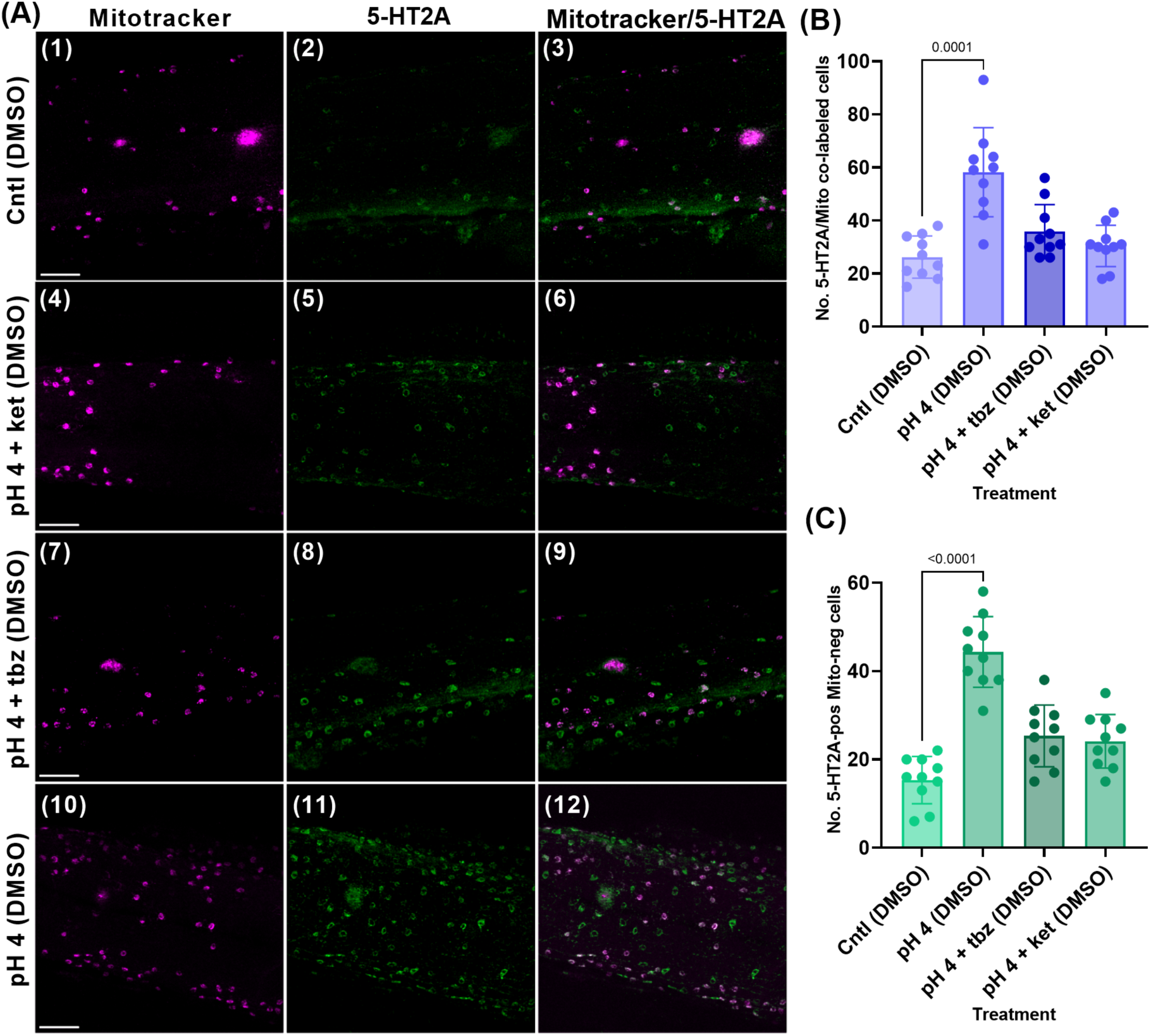
Treatment with ketanserin or tetrabenazine inhibited the increase in ionocyte number in acid-treated fish. (A) Representative confocal images of Mitotracker (magenta) labeling (1,4,7,10), 5-HT2A (green) labeling (2,5,8,11) and Mitotracker/5-HT2A co-labeling (white) (3,6,9,12) on the trunk of zebrafish larvae at 6 days postfertilization (dpf) under: control medium with 0.1% DMSO (1-3), pH 4 medium supplemented with 100 𝜇M ketanserin and 0.1% DMSO (4-6), pH 4 medium supplemented with 100 𝜇M tetrabenazine and 0.1% DMSO (4-6) and pH 4 medium with 0.1% DMSO (7-9) conditions. Fish were placed in the corresponding medium at approximately 1 h postfertilization and treated for 6 days. 10 fish from each of the four treatment groups were treated and processed for immunohistochemistry. Scale bars in A (1,4,7,10) are 50 𝜇m and apply to all panels in that row. (B) The number of ionocytes co-labeled with Mitotracker and 5-HT2A. (C) The number of ionocytes labeled with 5-HT2A only. Data analyzed using a Kruskal-Wallis test (two-tailed) with Dunn’s multiple comparison test. p values are shown on the graph for groups that are significantly different (p<0.05, N=10 for each group). Controls for panels (B) and (C) were performed at the same time as those for Fig. 7 and are derived from the same data set. Average values represented as mean±s.d. 5-HT, serotonin; 5-HT2A, serotonin 2A receptor; ket, ketanserin; tbz, tetrabenazine; DMSO, dimethyl sulfoxide; Cntl, control; Mito, Mitotracker; pos, positive; neg, negative.

Having demonstrated that 5-HT2A receptors are involved in the proliferation of ionocytes in an acidic environment, we also tested whether there was an endogenous source of 5-HT, such as from nearby cutaneous NECs, that may be activating this process. We used tetrabenazine to prevent the release of 5-HT from potential nearby sources. Tetrabenazine disrupts the loading and storage of monoamines through inhibition of vmat2 and has been shown to inhibit 5-HT-mediated signaling in gill NECs in zebrafish (Reed et al., 2025). When exposed to acidic medium supplemented with tetrabenazine, larvae demonstrated no significant increase (p>0.05, N=10) in ionocyte number compared to the negative control (Fig. 9B,C).

The number of ionocytes remained significantly upregulated in acid-treated fish in the absence of ketanserin and tetrabenazine for cells co-labelled with 5-HT2A and Mitotracker, and cells positive for 5-HT2A only (p=0.0001 and p<0.0001 respectively, N=10) (Fig. 9B,C).

## DISCUSSION

This study presents the novel finding that cutaneous ionocytes in developing zebrafish express 5-HT2A receptors. We have provided evidence that the increase in ionocyte number following acid acclimation is mediated, at least in part, by an endogenous source of 5-HT and activation of 5-HT2A. A proposed scheme for a pathway through which 5-HT mediates ionocyte proliferation is discussed in the following sections and is summarized in Figure 10.

**Figure 10.**
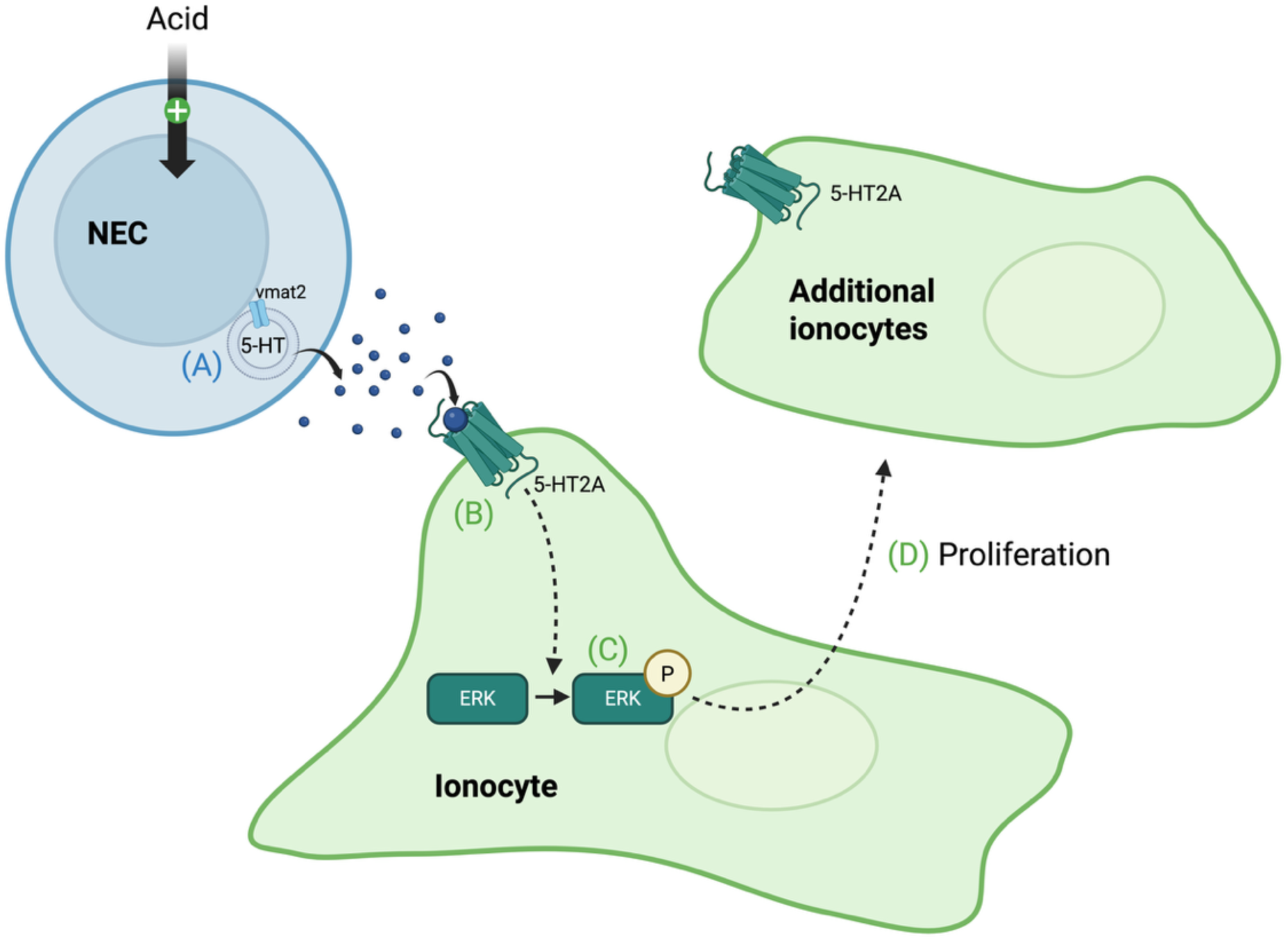
Model of a proposed pathway through which acid acclimation may lead to cutaneous ionocyte proliferation in zebrafish. (A) In a nearby cell, serotonin (5-HT) is loaded into secretory vesicles by the vesicular monoamine transporter 2 (vmat2) and is stored until released by acid stimulation. The only known source of cutaneous 5-HT in developing zebrafish are chemosensitive neuroepithelial cells (NEC; Coccimiglio and Jonz, 2012). (B) 5-HT binds to and activates 5-HT2A receptors on ionocytes to initiate an intracellular signaling cascade. (C) The signaling cascade results in phosphorylation (P) of extracellular signal-related kinase (ERK), a regulator of cell division and differentiation. (D) Phosphorylated ERK leads to proliferation of new 5-HT2A-positive ionocytes. See Discussion for further details.

### Ionocytes contain 5-HT2A receptors

The immunohistochemical data from experiments using 𝛼5, ConA, and Mitotracker demonstrate constitutive expression of 5-HT2A receptors in NaR and HR cells, as well as in MRCs. Furthermore, given the near complete co-localization of 5-HT2A labeling with Mitotracker dye, it seems likely that all cutaneous ionocyte subtypes contain 5-HT2A in zebrafish larvae until at least 7 dpf. Some cells reported in the present study were positive for 5-HT2A but did not take up Mitotracker. Despite not being co-labeled with this common ionocyte marker, these cells were likely ionocytes based on their morphology, distribution and response to acidic stimuli. This population represented approximately 24% of 5-HT2A-positive ionocytes and may correspond to SLC26 and KS ionocytes, which have not yet been demonstrated to stain with Mitotracker. Alternatively, these cells may be MRCs. It is important to note that Mitotracker dyes have been shown to be inconsistent in some cases (Neikirk et al., 2023) and may underestimate the labeling of cells that are rich in mitochondria, particularly in whole-mount preparations, as in the present study. We continued to differentiate between 5-HT2A-positive ionocytes, with and without Mitotracker labeling, and in each case ionocytes responded the same way to stimuli.

### Activation of 5-HT2A receptors mediates an increase in ionocyte number

The effects of 5-HT on ionocyte activity have not previously been recognized. In one study, Kumai et al. (2012) demonstrated that 5-HT had no effect on Na^+^ uptake in HR cells in zebrafish. We hypothesized that 5-HT2A receptors play a role in ionocyte proliferation. The increase in ionocyte number following exposure to 5-HT, and inhibition of this effect by pre-exposure to ketanserin, a 5-HT2A receptor antagonist, strongly suggest that this is the case.

This finding agrees with the literature demonstrating that 5-HT receptors are regulators of the cell cycle. A review by Azmitia (2001) highlights multiple examples of 5-HT2A and 5-HT1A receptors in regulation of cell proliferation, maturation, and apoptosis. Similarly, Masson et al. (2012) discussed a mitogenic role for 5-HT2A receptors, highlighting induction of growth of smooth muscle cells and potentiation of the activity of growth factors. Furthermore, 5-HT, via 5-HT2A receptors, has known roles in facilitating proliferation of neurons in the developing neocortex (Xing et al., 2020), is known for its mitogenic effects on trophoblast cells (Fecteau and Eiler, 2001; Oufkir et al., 2010; Sonier et al., 2005), and has a proliferative effect on rat renal mesangial cells (Takuwa et al., 1989).

The signaling pathways through which 5-HT2A receptors lead to cell proliferation are complex, but many examples implicate phosphorylation of ERK (Göőz et al., 2006; Greene et al., 2000; Grewal et al., 1999; Masson et al., 2012; Oufkir et al., 2010). We found that approximately 74% of 5-HT2A-positive ionocytes contained p-ERK in 5-HT-treated fish, and this effect was reduced to about 23% when fish were pre-treated with ketanserin. These results implicate phosphorylation of ERK as part of a signaling pathway through which 5-HT2A receptor activation increases ionocyte number in zebrafish larvae. Furthermore, the presence of p-ERK in ionocytes stimulated with 5-HT provides direct evidence for the activation of a pathway in 5-HT2A-positive ionocytes.

### 5-HT2A receptor activation occurs in response to osmotic stress

Given that 5-HT levels are known to increase in the brain (Boeck et al., 1996; Sreelekshmi et al., 2022) and gills (Mustafayev and Mekhtiev, 2008) of fishes exposed to osmotic stress, one of the aims of our study was to uncover whether 5-HT mediates the increase in ionocyte number under similar conditions. This hypothesis was supported by our demonstration that ionocyte proliferation, in response to an acidic environment, was inhibited by tetrabenazine and ketanserin, which deplete 5-HT stores and inhibit 5-HT2A, respectively. These findings demonstrate for the first time that 5-HT, via the 5-HT2A receptor, plays a role in ionocyte proliferation as zebrafish larvae acclimate to an acidic environment, and perhaps to other osmotic stressors. Interestingly, in juvenile carp, increasing dietary tryptophan (a precursor of 5-HT) has previously been demonstrated to result in increased salinity tolerance (Hoseini and Hosseini, 2010). It is not yet known whether ionocytes in carp contain 5-HT2A receptors, but future studies may determine whether ionocyte proliferation due to increased 5-HT production may have implications for the observed increase in tolerance to osmotic stress.

Our results suggest that release of endogenous 5-HT mediates the increase in ionocyte number in response to acid exposure. The source of 5-HT, however, remains to be elucidated. Chemosensitve NECs store 5-HT, express vmat2 and are found in close proximity to ionocytes in the skin and gills in zebrafish (Coccimiglio and Jonz, 2012; Dean et al., 2017; Jonz and Nurse, 2006; Pan et al., 2021). NECs are promising candidates for regulation of cutaneous ionocyte proliferation because they are the only known source of cutaneous 5-HT in developing zebrafish. In the gills, 5-HT depletion by tetrabenazine inhibits serotonergic signaling from NECs to postsynaptic neurons that express 5-HT3 receptors (Reed et al., 2025). In the present study, the inhibition of ionocyte proliferation by tetrabenazine suggests that 5-HT storage and release from NECs is an integral part of regulating ionocyte proliferation. Moreover, gill NECs express acid-sensitive TASK-2 ion channels (Peña-Münzenmayer et al., 2014) and have been shown to produce excitatory signals in response to extracellular H^+^ (Abdallah et al., 2015), a response that leads to neurosecretion of 5-HT from NECs (Reed et al., 2025). Whether skin NECs release 5-HT in response to acid exposure remains an important question for future investigation.

### Implications and conclusions

As we continue to gain insight into the complex mechanisms that underly ionocyte proliferation in response to osmotic stress, identification of the cell type that releases 5-HT to activate this pathway will be paramount, as will the identification of the specific source of increased ionocyte number. It has been suggested by Horng et al. (2009) that additional HR cells in pH 4-exposed fish may differentiate from proliferating p63-positive epithelial stem cells or other ionocyte precursors. Should this be the case for all ionocytes, the signaling cascade downstream of p-ERK may involve spatial propagation of ERK signals from activated ionocytes to ionocyte precursors, leading to regulation of proliferation and differentiation of ionocyte precursors to elicit the increase in ionocyte number. There is precedence for such a mechanism in the mammalian epidermis with implications in the regulation of cell density during development and wound healing (Aoki et al., 2013; Hiratsuka et al., 2015; Hiratsuka et al., 2020). Alternatively, it is possible that the signaling cascade initiated by 5-HT2A results in direct proliferation of ionocytes. To differentiate between these possibilities, more studies on the cellular origins of ionocytes in acid-acclimated zebrafish larvae, and expression profiles of ionocyte precursors, would be required.

Furthermore, identification of the remaining molecular players upstream and downstream of p-ERK will allow further comparisons to be drawn between ionocytes and other cell types in vertebrates. As our understanding of this pathway expands, this research may lead to implications for the use of zebrafish as a model organism, should the pathway continue to present parallels with similar cell types in mammals. Finally, investigation of the presence of 5-HT2A and the importance of this pathway in gill ionocytes of adult zebrafish, and ionocytes of other fish species is warranted. Should the same mechanism be present in ionocytes in other fish species, this research may have applications for fish rearing in osmotically stressful environments.

## Acknowledgements

W.M.M. was supported by an Undergraduate Student Research Award from the Natural Sciences and Engineering Research Council of Canada (NSERC).

## Author Contributions

Conceptualization: W.M.M., M.G.J.; Methodology: W.M.M., M.G.J.; Software: M.G.J.; Validation: W.M.M., M.G.J.; Formal analysis: W.M.M., M.G.J.; Investigation: W.M.M., M.G.J.; Resources: M.G.J.; Data curation: W.M.M., M.G.J.; Writing - original draft: W.M.M.; Writing - review & editing: W.M.M., M.G.J.; Visualization: W.M.M., M.G.J.; Supervision: M.G.J.; Project administration: M.G.J.; Funding acquisition: M.G.J.

## Data and resource availability

All relevant data and details of resources can be found within the article and its supplementary information.

---

**Supplementary Figure 1.**
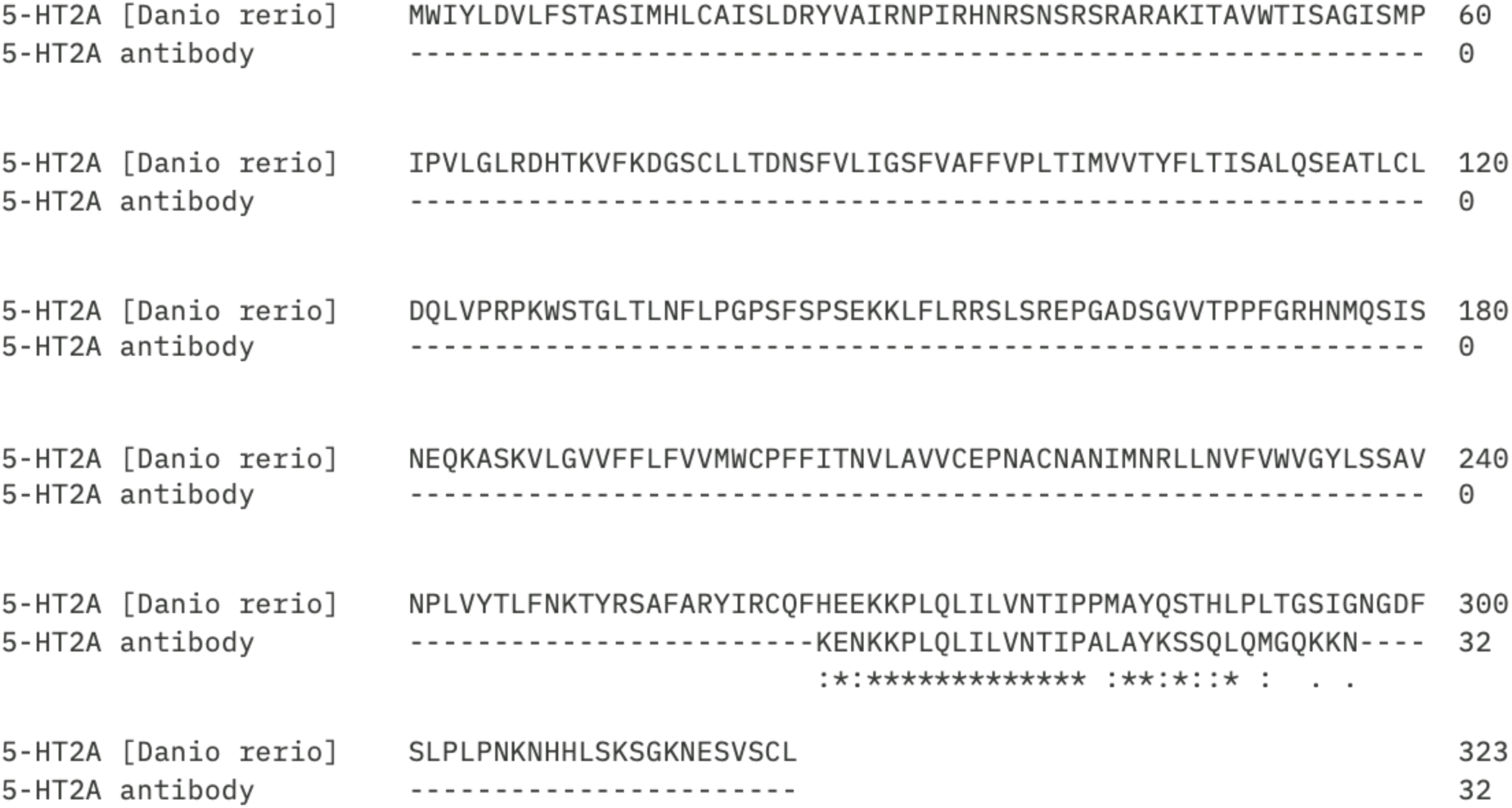
Sequence alignment for the antigen of the 5-HT2A antibody compared to the zebrafish 5-HT2A receptor. Sequence alignment from ClustalOmega (URL: https://www.ebi.ac.uk/jdispatcher/msa/clustalo, accessed Oct. 18, 2025) for the antigen of the 5-HT2A antibody (manufacturer specifications) with the sequence of the zebrafish (*Danio rerio*) 5-HT2A receptor (NCBI reference sequence XP_068079731.1) obtained from NCBI (URL: https://www.ncbi.nlm.nih.gov/, accessed Oct. 18, 2025). (.) represents semi-conservative substitutions, (:) conserved substitutions and (*) fully conserved residues.

**Supplementary Figure 2.**
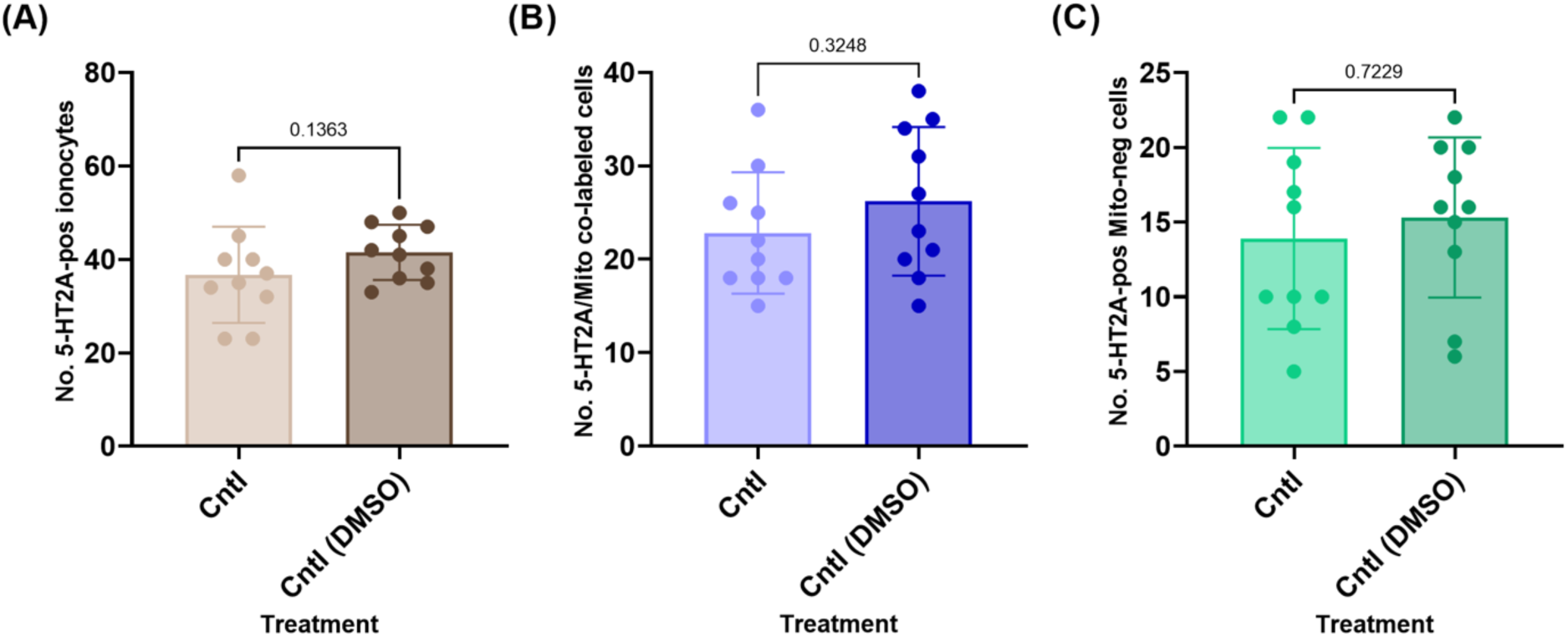
DMSO did not significantly change the number of 5-HT2A-positive ionocytes. The addition of 0.1% DMSO did not significantly alter (p>0.05, N=10) (A) the total number of 5-HT2A positive ionocytes, (B) the number of ionocytes co-labeled with 5-HT2A and mitotracker, or (C) the number of 5-HT2A-positive ionocytes that were Mitotracker-negative. Fish treated without DMSO (Control, Cntrl) and with DMSO were exposed and processed for immunohistochemistry. Data analyzed using a Mann-Whitney test (two-tailed) (p<0.05, N=10 for each group). All p values are shown on the graphs. Average values represented as mean±s.d. 5-HT, serotonin; 5-HT2A, serotonin 2A receptor; DMSO, dimethyl sulfoxide; pos, positive.

